# Effects of mind-wandering on cognitive and neural processes: Identifying specific impairments in adults with attention-deficit/hyperactivity disorder

**DOI:** 10.1101/2025.09.19.677328

**Authors:** Gabriela Horáková, Sébastien Weibel, Ugo Tissot, Bich-Thuy Pham, Madalina Elena Costache, Anne Bonnefond

## Abstract

Attention-deficit/hyperactivity disorder (ADHD) is a common neurodevelopmental disorder characterized by symptoms of inattention and/or impulsivity/hyperactivity. Although not a core symptom, excessive mind-wandering (MW) is commonly associated with the disorder and may contribute to symptoms as well as performance deficits observed in these patients. The main aim behind the present study is to better characterize MW and its impact on performance measures as well as on neural activity in adults with ADHD.

Twenty-eight medication-naïve ADHD patients and 28 healthy controls were matched on sex, age, and level of education; they completed a prolonged Go/No-Go sustained attention task including thought-probes to detect episodes of MW. Analysis of EEG data focused on P100, P3b, and CRN event-related potentials reflecting early visual, attention allocation, and performance monitoring processes, respectively.

MW, particularly spontaneous MW, was more frequent in patients and linked to a more severe ADHD symptomatology. A differentiated negative impact of MW on performance was noted in both groups: speed and variability of patients’ reaction times and accuracy in controls were affected. If the same perceptual (P100) and attentional (P3b) impairments were evidenced in both groups during MW, then the impairment of the performance monitoring (CRN) process was exclusively highlighted in ADHD patients (as revealed by a state × group interaction).Taken together, these results extend the understanding of the cognitive and neural processes associated with increased MW in patients and suggest performance monitoring as a potential key mechanism underlying specific MW-related impairments in ADHD.

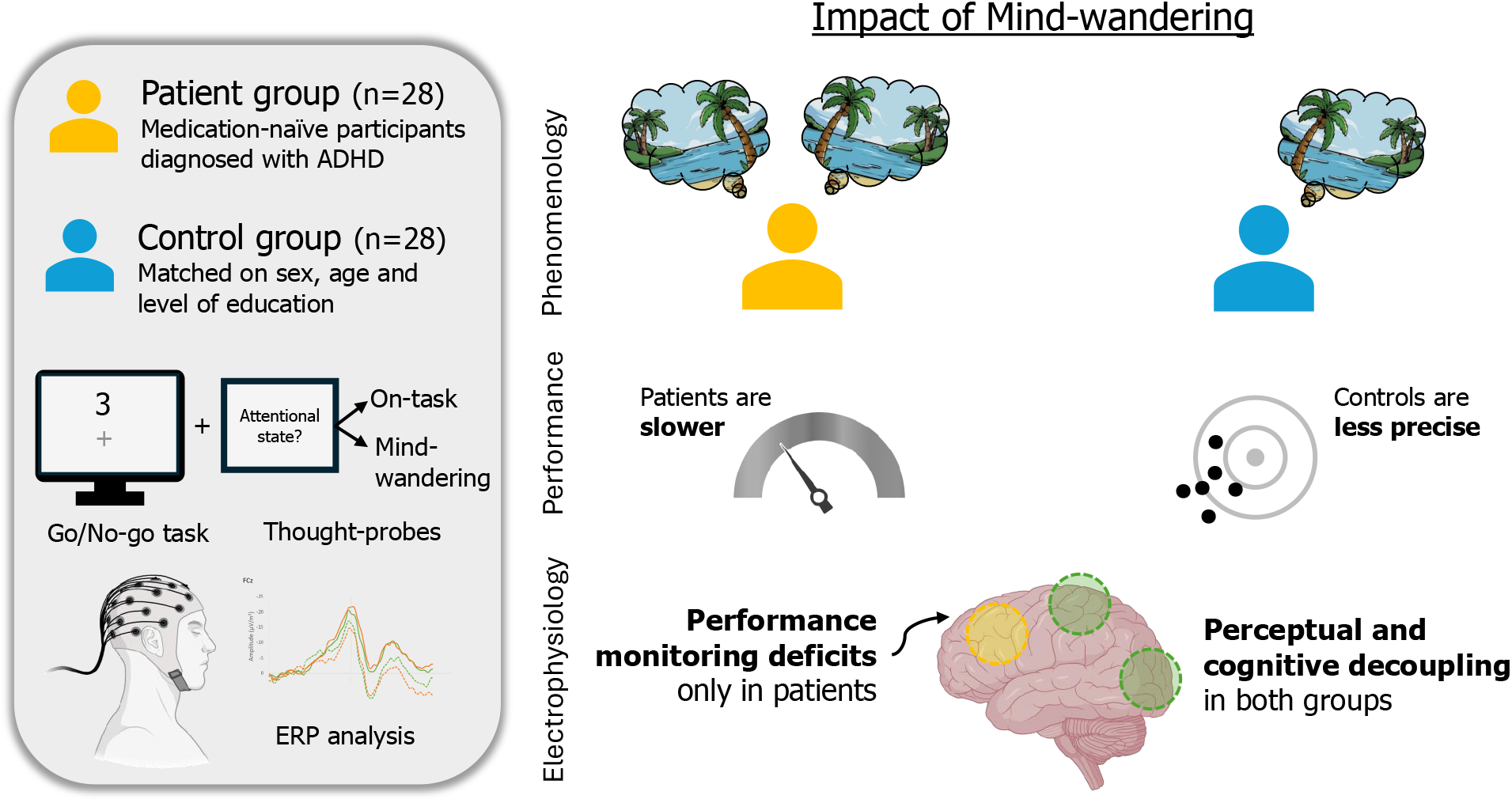

## 1. Introduction

Attention-deficit/hyperactivity disorder (ADHD) is a common neurodevelopmental disorder characterized by persistent symptoms of inattention and/or impulsivity/hyperactivity that disrupt normal functioning or developmental progress (American Psychiatric Association, 2013); its estimated prevalence is 3%–5% of the adult population (Fayyad et al., 2007; Thomas et al., 2015). Although not a core symptom, excessive mind-wandering (MW)—a shift of attention from a task toward internal thoughts (Smallwood and Schooler, 2006)—is a commonly associated feature of ADHD; however, most existing research has relied on trait-level questionnaires, leaving its real-time phenomenology and underlying neural processes largely unexplored. MW has been linked to reduced deactivation of the default mode network (DMN) and disrupted interactions with the frontoparietal (FPN) and salience networks (SN) (Christoff et al., 2009; Fox et al., 2015; Stawarczyk and D’Argembeau, 2015). Given that similar DMN dysregulation is observed in patients with ADHD (Gao et al., 2019; Sutcubasi et al., 2020), a recent hypothesis proposes that DMN hyper-activation may drive excessive MW in ADHD, thereby contributing to inattention symptoms and performance impairments (Bozhilova et al., 2018). Supporting this hypothesis, evidence demonstrates that the frequency of MW, particularly spontaneous or unintentional MW (MW-S), is positively correlated with the severity of all ADHD symptoms (Arabacı and Parris, 2018; Bozhilova et al., 2021a; Madiouni et al., 2020) and is associated with performance impairments (Alperin et al., 2021; Bozhilova et al., 2021a; Gau et al., 2022). Additionally, findings from studies that use the thought-probe method, which involves stopping participants during a task and asking them where their attention was directed, indicate that healthy controls during episodes of MW exhibit similar performance deficits (Stawarczyk et al., 2014, 2011) as those typically described in individuals with ADHD: slower, more variable, and less accurate responses (Gmehlin et al., 2016; Johnson et al., 2007; Kofler et al., 2013). However, studies that use clinical samples and the thought-probe method are rare and have revealed mixed results at the behavioral level. Indeed, some have highlighted similar MW-related impairments across both patient and control groups (Alperin et al., 2021; Gau et al., 2022; Madiouni et al., 2020), while others report that altered accuracy and speed was exacerbated by MW in ADHD patients only (Bozhilova et al., 2021a). These disparities may be linked to the diversity of methodologies used, such as the framing of the thought-probe and response options, the nature of cognitive tasks, or even the length of the analyzed window.

According to the decoupling phenomenon, broadly observed in healthy controls, MW is associated with lower P100 and P3b amplitudes, two event-related potentials (ERP) that reflect early sensory processing and attention allocation, respectively (Kam et al., 2022). One study also found altered performance monitoring processes during episodes of MW, as revealed by lower amplitudes of feedback error-related negativity (fERN) (Kam et al., 2012). Concurrently, overall attention allocation deficits are characteristic of ADHD patients (Szuromi et al., 2011) and recent studies also show impaired performance monitoring in these patients (Ehlis et al., 2018; Marquardt et al., 2018). To the best of our knowledge, only 2 studies took into account patients’ attentional state when investigating only the P100 and P3b components (Bozhilova et al., 2021a, 2020). Notably, the results reveal a consistent pattern of perceptual decoupling in individuals with ADHD regardless of attentional state, as evidenced by lower P100 amplitudes compared to healthy controls during both MW and on-task (OT; i.e., task-related thoughts) episodes (Bozhilova et al., 2021a). Regarding attentional processing deficits in ADHD, which are assessed through the P3b component, the findings indicate that the reduced amplitude of this ERP component is only partially linked to a MW tendency and symptomatology (Bozhilova et al., 2020). Instead, task conditions appear to be the primary influencing factor, rather than the attentional state (Bozhilova et al., 2021a).

To precisely investigate MW and its effects in ADHD, we combined a thought-probe method with EEG recordings during a prolonged Go/No-Go sustained-attention-to-response task (SART). Our first aim with the present study is to better characterize the phenomenology of MW in patients with ADHD by comparing the frequency of spontaneous (MW-S) and deliberate (deliberate (MW-D) mind-wandering. To further address this aim, we also examined how both types of MW relate to symptom severity and other subjective states, such as motivation, boredom, and fatigue. Our second aim is to investigate the impact of MW on performance measures of accuracy (commission errors) and speed (reaction times and reaction time variability) as well as the impact of MW on neural activity. Consequently, we focused on the P100 and P3b—the ERP correlates of early visual and attentional processing—as well as correct response negativity (CRN), an ERP component reflecting performance monitoring processes and provides insight into the activity of the medial prefrontal cortex (mPFC).

## 2. Methods and materials

### 2.1 Subjects

Our sample (*N* = 56) comprised a patient group of medication-naïve participants with ADHD (*n* = 28) and a control group (*n* = 28). Participants, aged 18-59 years (mean and standard deviation [*SD*] 32.7 ± 11.4 years), were matched on sex, age, and level of education (Table 1). Each subject was required to meet the inclusion criteria: the absence of neurological or psychiatric comorbidities, no history of psychostimulant medication treatment, and no record of head trauma with loss of consciousness. Participants with ADHD were diagnosed by a psychiatrist according to the DSM-5 criteria (American Psychiatric Association, 2013) prior to the study. All subjects completed the French version of the adult ADHD self-report scale v1.1 (ASRS) (Adler et al., 2006) to assess adult ADHD symptomatology; this 18-item questionnaire is used to evaluate ADHD symptoms over the past 6 months using a 5-point Likert scale ranging from “Never = 0” to “Very often = 4.” Separate scores for inattentive symptoms and impulsivity/hyperactivity were calculated for each subject based on the mean score of the nine items that corresponded to each dimension. This study was approved by the clinical research and innovation department of the University Hospitals of Strasbourg (API 2017 HUS N° 7248); all participants provided their consent and were financially compensated for their participation.

**Table 1.**
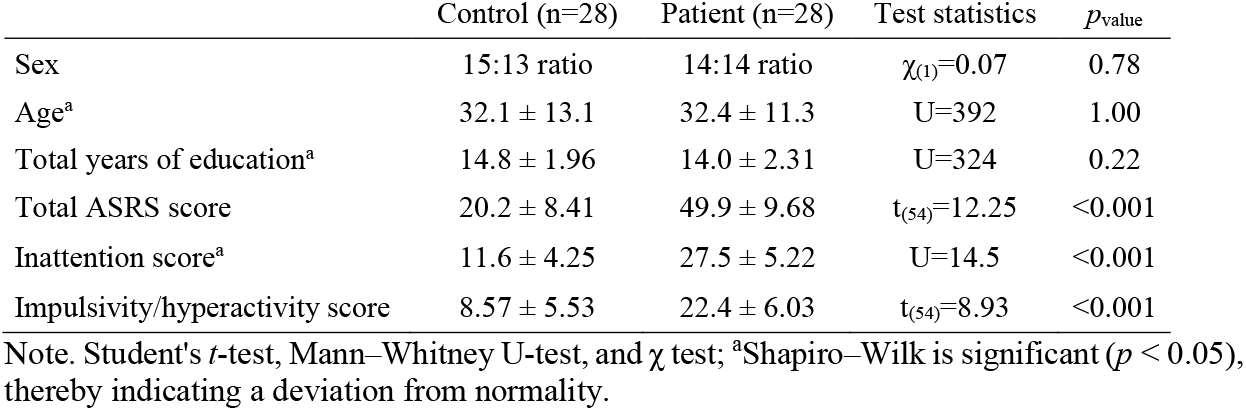
Demographic characteristics and ASRS scores for the control and patient groups (mean ± standard deviation)

| | Control (n=28) | Patient (n=28) | Test statistics | $p_{\text{value}}$ |
| --- | --- | --- | --- | --- |
| Sex | 15:13 ratio | 14:14 ratio | $\chi_{(1)}=0.07$ | 0.78 |
| Age<sup>a</sup> | $32.1 \pm 13.1$ | $32.4 \pm 11.3$ | U=392 | 1.00 |
| Total years of education<sup>a</sup> | $14.8 \pm 1.96$ | $14.0 \pm 2.31$ | U=324 | 0.22 |
| Total ASRS score | $20.2 \pm 8.41$ | $49.9 \pm 9.68$ | $t_{(54)}=12.25$ | <0.001 |
| Inattention score<sup>a</sup> | $11.6 \pm 4.25$ | $27.5 \pm 5.22$ | U=14.5 | <0.001 |
| Impulsivity/hyperactivity score | $8.57 \pm 5.53$ | $22.4 \pm 6.03$ | $t_{(54)}=8.93$ | <0.001 |

Note. Student's $t$ -test, Mann–Whitney U-test, and $\chi$ test; <sup>a</sup>Shapiro–Wilk is significant ( $p < 0.05$ ), thereby indicating a deviation from normality.

### 2.2 The SART

Participants completed the SART (Robertson et al., 1997), a computerized Go/No-Go task in which single-digit numbers from 1 to 9 were randomly presented with an equivalent probability of appearance (1/9). Participants were instructed to press the response button whenever a digit appeared on the screen (Go trial), except for the digit 3, which required no response (No-Go trial). They were encouraged to be quick and accurate. Each digit appeared in the center of the screen for 167 ms, which was followed by an inter-stimulus interval that varied between 1.5 s and 2.5 s (1500 ms, 1700 ms, 2100 ms, 2300 ms, and 2500 ms) that was reinitialized after each response. The digits varied in size (100, 120, 140, 160, or 180) in order to enhance the demands for processing the numerical value and were presented in Arial font. They were displayed in black type and were positioned 0.25° above a central yellow fixation cross on a gray background; the computer screen was a 17-inch CRT monitor. A total of 810 trials (720 Go trials and 90 No-Go trials) were divided into 15 blocks of either 27, 54, or 81 trials. The whole task required approximately 40 minutes per participant. At the end of each block, a thought-probe of four questions was presented on the screen. The first question asked participants about their attentional state immediately before the probe. Three response options were possible: 1) on-task (OT); 2) spontaneous mind-wandering (MW-S) and 3) deliberate mind-wandering (MW-D). Participants were instructed to select OT if their thoughts or attention were mainly engaged with the task, MW-S if their thoughts or attention drifted from the task unintentionally, and MW-D if they intentionally allowed their thoughts or attention to drift from the task. The frequency of each attentional state (OT, MW-S, MW-D) was calculated over the entire task as a percentage of the number of probes in which participants reported a specific attentional state of the 15 total probes. The subsequent three questions inquired regarding participants’ level of motivation, boredom, and fatigue (subjective states) by positioning on analog scales that ranged from “Lowest level = 0” to “Highest level = 100.” Task performance was assessed over a 20-trial pre-probe window using the following measures: the number of commission errors (CE), i.e. the number of errors in No-Go trials, the reaction time (RT) on Go trials in seconds, the standard deviation of RT (*SD*-RT) in seconds, and the efficiency score (ES), i.e. the number of correct withholds divided by the mean RT. To account for inter-individual differences in MW rate, the performance measures were pondered by the subject’s total MW frequency. Before beginning the SART, an explanation was provided to each participant to ensure that the instructions were understood, and a 3-minute training session was conducted.

### 2.3 EEG recording

The EEG was recorded with the BioSemi ActiveTwo System (Amsterdam, Netherlands) using a montage of 64 active electrodes, following the international 10-20 standard system. Two additional electrodes—a common mode sense (CMS) active electrode and a driven right leg (DRL) passive electrode placed on the nape—served mainly as reference and ground electrodes. The signal was acquired at a sampling rate of 2048 Hz. The electrooculography (EOG) was recorded by two electrodes placed at the outer canthi of the eyes and two electrodes placed above and below the right eye. Two supplementary reference electrodes were attached to the earlobes.

### 2.4 EEG preprocessing

The raw EEG data was preprocessed using custom scripts on Python v.3.11 via MNE library (version 1.7) (Gramfort et al., 2013). Bad channels were identified using PyPrep (Appelhoff et al., 2025; Bigdely-Shamlo et al., 2015) default criteria, such as correlation with other channels, deviation from the mean, high-frequency noise, flat signals, signal-to-noise ratio (SNR) and were further confirmed through visual inspection. A high-pass filter of 0.2 Hz and a low-pass filter of 30 Hz were applied, and the signal was re-referenced to the averaged signal of both earlobes. In order to eliminate ocular artifact-related components, a semiautomatic independent component analysis (ICA), using FastICA algorithm, was performed. To remove low-frequency drifts and facilitate ICA convergence, a high-pass filter at 1 Hz was applied. Components that explained at least 99% of the total variance were selected using a fixed seed for reproducibility. Components were preselected based on correlations (.70) with EOG channels and through qualitative inspection. The preselected bad channels were then interpolated (3.4 channels, on average, per subject).

### 2.5 ERP analysis

The segmentation was performed using BrainVision Analyzer 2.1 software (Brain Products GmbH, Germany). The signal was segmented into 54 s epochs (approximately 20 trials, the same window that was used for the performance analysis) prior to each probe and characterized as either OT or MW based on the subject’s answer. Stimulus-locked ERPs were epoched from −200 ms to 700 ms around Go and No-Go stimuli and the response-related ERP was epoched from −400 ms to 400 ms around responses on Go trials. All epochs were visually inspected, and artifacted segments were semi-automatically rejected based on the following criteria: maximal allowed voltage step of 50 µ*V*/ms and maximal absolute difference of 150 µ*V*. On average, 3.55 ± 15.59 trials per subject were removed. Current source density (CSD) transformation (Perrin et al., 1987) was applied to reduce the volume conduction effects; parameters were used (stiffness *m* = 4, λ = 10^−5^). Stimulus-locked ERPs were baseline corrected from −200 ms to 0 ms; response-related ERPs were baseline corrected from −400 ms to −200 ms. The P100 was defined as a positive peak; maximum amplitude (µ*V*/m^2^) over the Oz and POz electrodes within 100 ms to 200 ms after stimulus onset. The P3b was defined as the mean activity (µ*V*/m^2^) between 300 ms and 550 ms after stimulus onset, which was extracted over Pz and CPz electrodes. For the CRN, the most negative peak amplitude (µ*V*/m^2^)—between −30 ms and 100 ms—was extracted at the Fz and FCz electrodes, which was where the peak was the most apparent.

### 2.6 Statistical analysis

All statistical analyses were conducted with jamovi (version 2.3.24) computer software (The jamovi project, 2025). The condition of normality of the data distribution was systematically tested. If the data was normally distributed, then parametric tests were applied; otherwise, the data was either transformed using a logarithmic transformation or nonparametric tests were applied. Independent samples *t*-tests were used to compare MW-S and MW-D frequencies between groups, and correlation analyses were performed to examine the relationships between each type of MW frequency and ASRS scores. Predictors of MW-S and separately MW-D (dependent variables) were specified by multiple linear regression analyses; we used the ASRS scores, the levels of subjective states (motivation, boredom and fatigue), and the group factor as possible predictors (independent variables. To identify the most parsimonious model, a backward elimination method was applied, using AIC and R^2^ values to guide model selection. Performance and ERP measures were subjected to an analysis of variance (ANOVA) including the within-subject factor of state (OT or MW) and the between-subject factor of group (control or patient). For ERP measures only, the within-subject factor electrodes (Oz or POz for the P100; Pz or CPz for the P3b, and Fz or FCz for the CRN) were added. Effect sizes were calculated using partial eta squared (η^2^*p*), and Tuckey post-hoc tests that compared different conditions were applied for all repeated measures ANOVAs. Additional analyses and results are available in the supplementary materials.

## 3. Results

### 3.1 Frequency and characteristics of the MW states

A student’s *t*-test performed on MW-S indicated a significantly higher frequency in patients (43%) than in controls (23%) (*t*(54) = 4.24, *p* < 0.001). A Mann–Whitney U-test performed on MW-D indicated a trend toward a higher frequency in patients (19%) than in controls (9%) (U = 173, *p* = 0.08). Spearman correlation analyses revealed that the MW-S frequency was positively correlated with inattention (*rs* = 0.40, *p* < 0.05), impulsivity/hyperactivity (*rs* = 0.37, *p* < 0.001), and total ASRS scores (*rs* = 0.41, *p* < 0.01). MW-D frequency was positively correlated only with impulsivity/hyperactivity (*r*_*s*_ = 0.38, *p* < 0.001) and total ASRS scores (*rs* = 0.29, *p* < 0.05). Multiple linear regressions revealed that level of boredom (β = 0.22, *t* = 2.41, *p* < 0.05) and group (P-T = β = 16.91, *t* = 2.41, *p* < 0.01) were predictors of MW-S (*R*^2^ = 0.32, *F*_(2,53)_ = 12.7, *p* < 0.001), whereas the level of boredom (β = 0.20, *t* = 2.92, *p* < 0.01) and impulsivity/hyperactivity ASRS score (β = 0.56, *t* = 2.82, *p* < 0.01) were predictors of MW-D (*R*^2^ = 0.28, *F*_(2,53)_ = 10.4, *p* < 0.001).

### 3.2 Impact of MW on performance and ERP measures

A total of 32% of participants (8 patients and 10 controls) did not report any MW-D episodes. Distinguishing between the two MW types was no longer possible due to the insufficient number of MW-D trials for performance and ERP analyses. Consequently, MW-S and MW-D were pooled and referred to collectively as MW in the subsequent analyses. Mean ± *SD* according to state (OT or MW) and group (control or patient) are reported in Table 2. Moreover, two participants (2 patients) were never OT and 3 participants (1 patient and 2 controls) never reported either type of MW. Missing EEG data for these 5 subjects who remained in the same attentional state throughout the entire task, were replaced by the mean ERPs of their respective group.

**Table 2.**
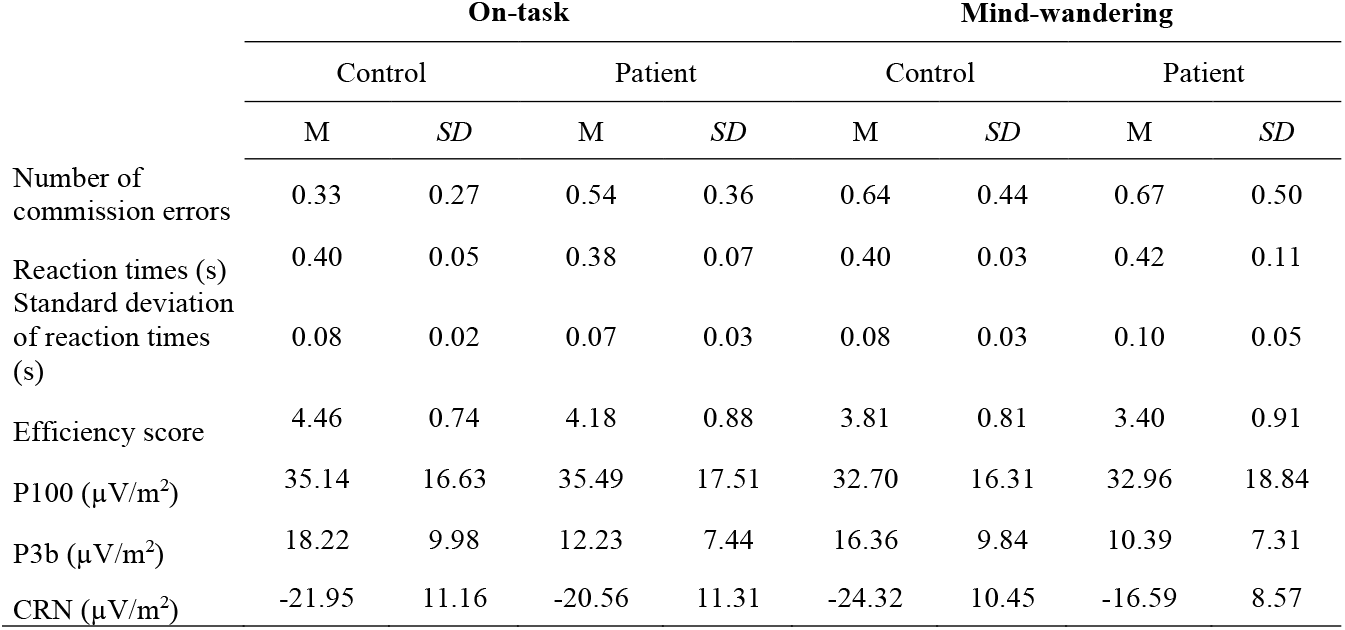
Performance and event-related potential (ERP) measures according to state (on-task or mind-wandering) and group (control or patient) (mean ± standard deviation).

|  | On-task |  |  |  | Mind-wandering |  |  |  |
| --- | --- | --- | --- | --- | --- | --- | --- | --- |
|  | Control |  | Patient |  | Control |  | Patient |  |
|  | M | SD | M | SD | M | SD | M | SD |
| Number of commission errors | 0.33 | 0.27 | 0.54 | 0.36 | 0.64 | 0.44 | 0.67 | 0.50 |
| Reaction times (s) | 0.40 | 0.05 | 0.38 | 0.07 | 0.40 | 0.03 | 0.42 | 0.11 |
| Standard deviation of reaction times (s) | 0.08 | 0.02 | 0.07 | 0.03 | 0.08 | 0.03 | 0.10 | 0.05 |
| Efficiency score | 4.46 | 0.74 | 4.18 | 0.88 | 3.81 | 0.81 | 3.40 | 0.91 |
| P100 ($\mu\text{V}/\text{m}^2$) | 35.14 | 16.63 | 35.49 | 17.51 | 32.70 | 16.31 | 32.96 | 18.84 |
| P3b ($\mu\text{V}/\text{m}^2$) | 18.22 | 9.98 | 12.23 | 7.44 | 16.36 | 9.84 | 10.39 | 7.31 |
| CRN ($\mu\text{V}/\text{m}^2$) | -21.95 | 11.16 | -20.56 | 11.31 | -24.32 | 10.45 | -16.59 | 8.57 |

#### 3.2.1 Performance measures

The ANOVA performed on the number of CE revealed a state × group interaction: *F*_(1.40)_ = 4.56, *p* < 0.05, η^2^*p* = 0.10 (Figure 1.A). In controls, the number of CE was significantly higher during MW than OT episodes, whereas this number was similar between the two states in patients. The number of CE was also significantly higher in patients than in controls during OT episodes but was similar between the two groups during MW episodes. A main effect of state, *F*_(1.40)_ = 10.31, *p* < 0.01, η^2^*p* = 0.20, was observed, as was a main effect of group: *F*_(1.40)_ = 4.52, *p* < 0.05, η^2^*p* = 0.10. The number of CE was significantly higher during MW than OT episodes and higher in patients compared to controls.

**Figure 1.**
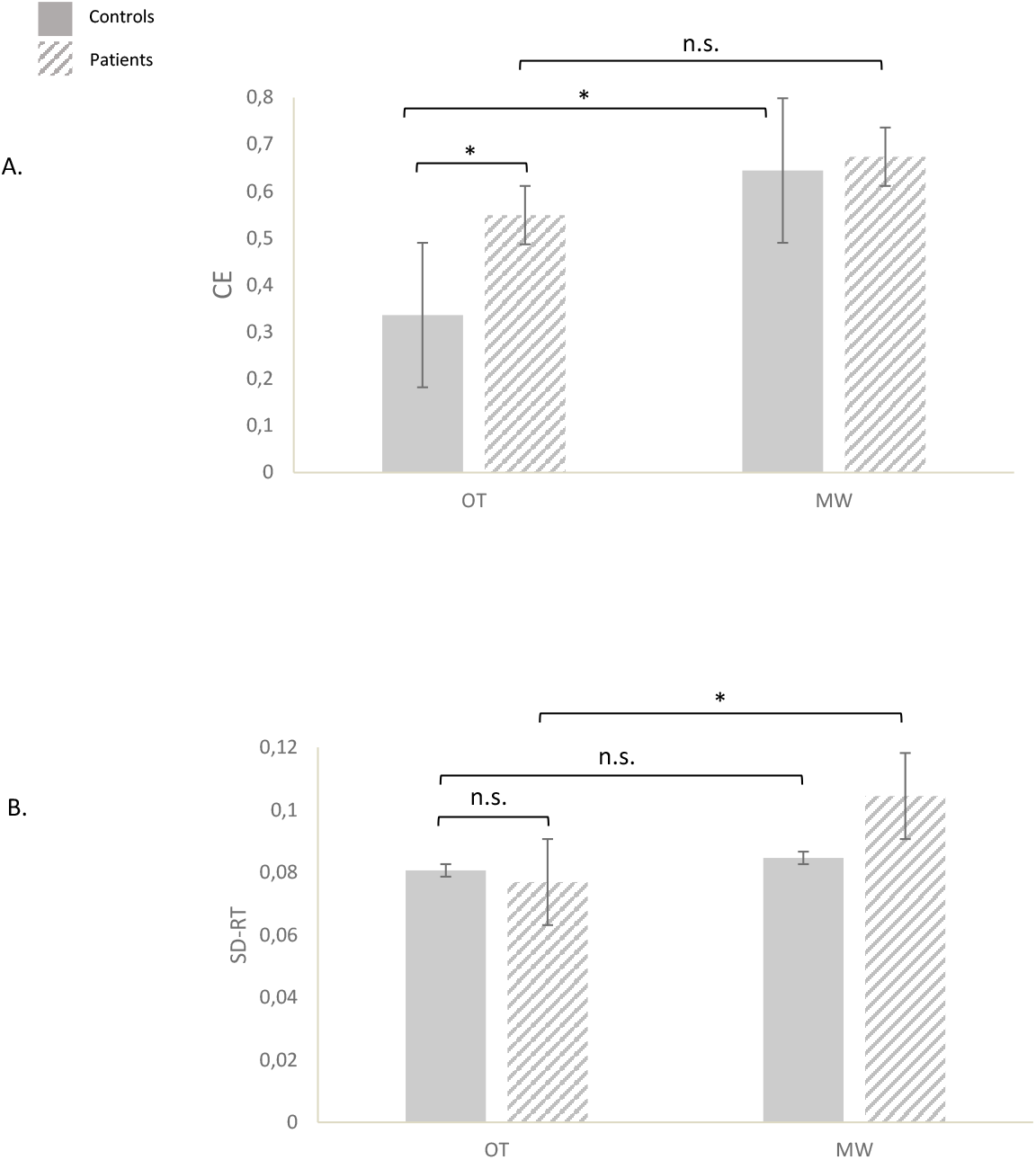
Performance measures. A. Number of commission errors (CE) in each group (control or patient) during on-task (OT) and mind-wandering (MW) episodes. B. Standard deviation of reaction times (*SD*-RT) in seconds in each group (control or patient) during on-task (OT) and mind-wandering (MW) episodes. \**p* < 0.05; n.s. = nonsignificant; error bars represent the standard error.

The ANOVA performed on RT revealed a significant state × group interaction: *F*_(1.49)_ = 4.12, *p* < 0.05, η^2^*p* = 0.07. In patients, RT was significantly longer during MW than OT episodes, whereas RT was similar between the two states in controls. A trend effect of state was also observed, *F*_(1.49)_ = 3.08, *p* =0.08, η^2^*p* = 0.05, which indicates that RT tended to be longer during MW than OT episodes. No effect of group was observed (*p* = 0.76).

The ANOVA performed on *SD*-RT revealed a significant state × group interaction: *F*_(1.49)_ = 4.90, *p* < 0.05, η^2^*p* = 0.09 (Figure 1.B). In patients, the *SD*-RT was significantly higher during MW than OT episodes, whereas this score was similar between the two states in controls. A main effect of s t a t e was also observed: *F*_(1.49)_ = 16.72, *p* < 0.01, η^2^*p* = 0.25. The *SD*-RT was significantly higher during MW than OT episodes. No main effect of group was observed (*p* = 0.63).

The ANOVA performed on the ES revealed no state × group interaction (*p* = 0.79). A main effect of state, *F*_(1.49)_ = 22.78, *p* < 0.001, η^2^*p* = 0.31, was observed, as was a main effect of group: *F*_(1.49)_ = 4.58, *p* < 0.05, η^2^*p* = 0.08. The ES was significantly lower during MW than OT episodes and lower in patients compared to controls.

To ensure that the larger 20-trial window did not affect our results, an ANOVA was performed on RT and *SD*-RT within a 10-trial window. This analysis revealed the same state × group interactions (p = 0.06) for both measures and revealed a main effect of state for RT (p = 0.08) and *SD*-RT (p < 0.01). An ANOVA on the number of CE and ES was not performed due to the lack of No-Go stimuli within the 10-trial window.

#### 3.2.2 ERP measures

Two patients and three controls were excluded from the EEG analysis due to poor data quality, which was characterized by numerous noisy channels in the same region that could not be interpolated. The EEG analysis was conducted on a final sample of 51 participants, which comprised 26 patients and 25 controls.

The ANOVA performed on the P100 amplitude revealed no state × group interaction (*p* = 0.96) (Figure 2.A). A main effect of state was observed: *F*_(1,49)_ = 6.10, *p* < 0.05, η^2^*p* = 0.11. The P100 amplitude was significantly lower during MW than OT episodes. No main effect of group (*p* = 0.94) was observed.

**Figure 2.**
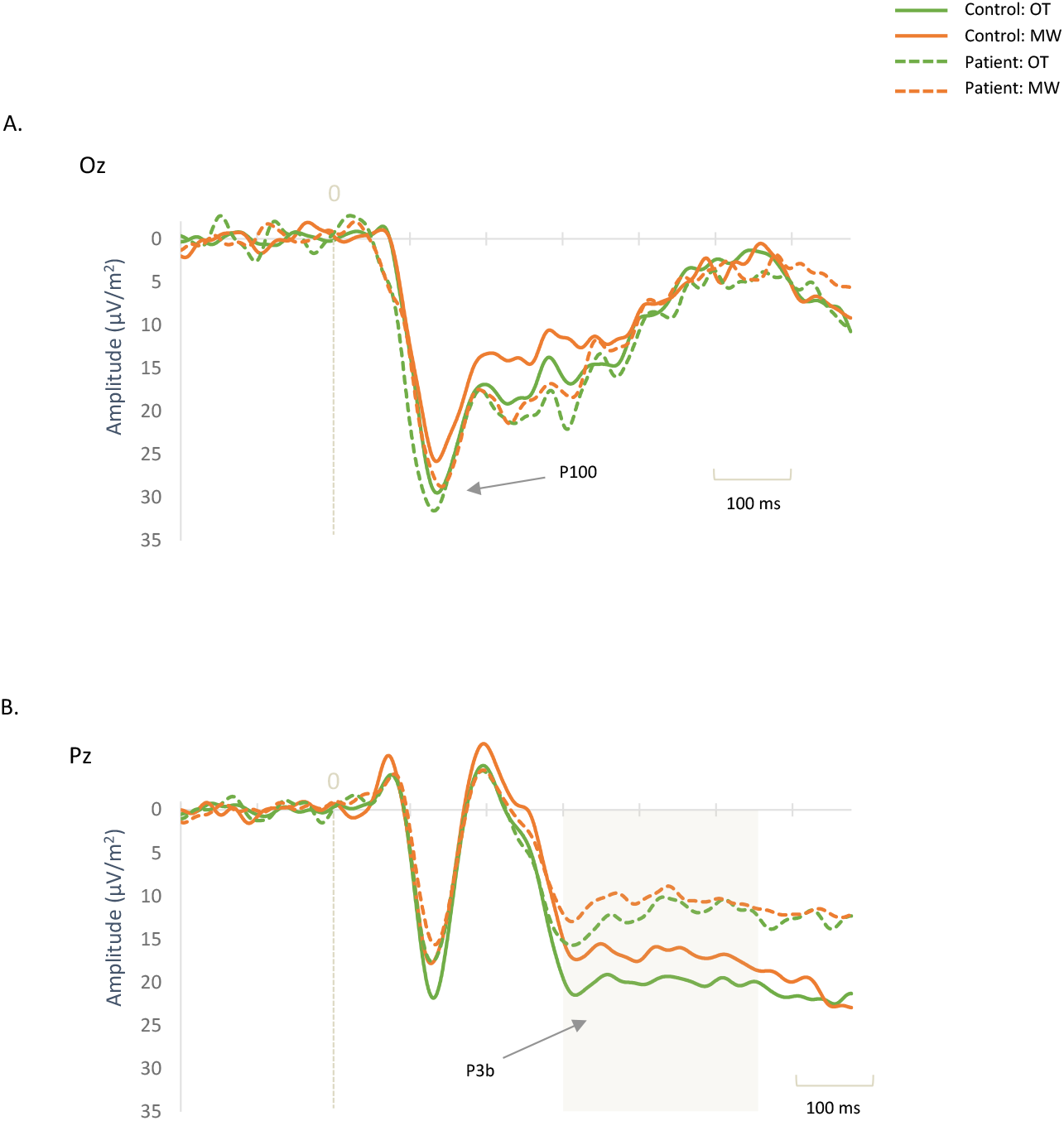
Grand average of a stimulus-locked event-related potential (ERP) according to state (on-task [OT] or mind-wandering [MW]) and group (control or patient). A. P100 at the Oz electrode; B. P3b at the Pz electrode. P100 and P3b amplitudes are significantly higher during OT than MW episodes (*p* < 0.05 and *p* < 0.01, respectively), and P3b amplitude is significantly higher in controls compared to patients (*p* < 0.01).

The ANOVA performed on the P3b mean activity revealed no state × group interaction (*p* = 0.98) (Figure 2.B). A main effect of state, *F*_(1,49)_ = 10.42, *p* < 0.01, η^2^*p* = 0.17, was observed, as was a main effect of group: *F*_(1,49)_ = 7.64, *p* < 0.01, η^2^*p* = 0.13. The P3b mean activity was significantly lower during MW than OT episodes and was lower in patients than in controls.

The ANOVA performed on the CRN amplitude revealed a significant state × group interaction: *F*_(1,49)_ = 9.43, *p* < 0.01, η^2^*p* = 0.16 (Figure 3). In patients, the CRN amplitude was significantly lower during MW than OT episodes, whereas it was similar between the two states in controls. The CRN was also significantly lower in patients than in controls during MW episodes, whereas it was similar between the two groups during OT episodes. A trend effect of group was also observed: *F*_(1,49)_ = 3.41, *p* = 0.07, η^2^*p* = 0.06. The CRN amplitude was significantly lower in patients than controls. No main effect of state was observed (*p* = 0.44).

**Figure 3.**
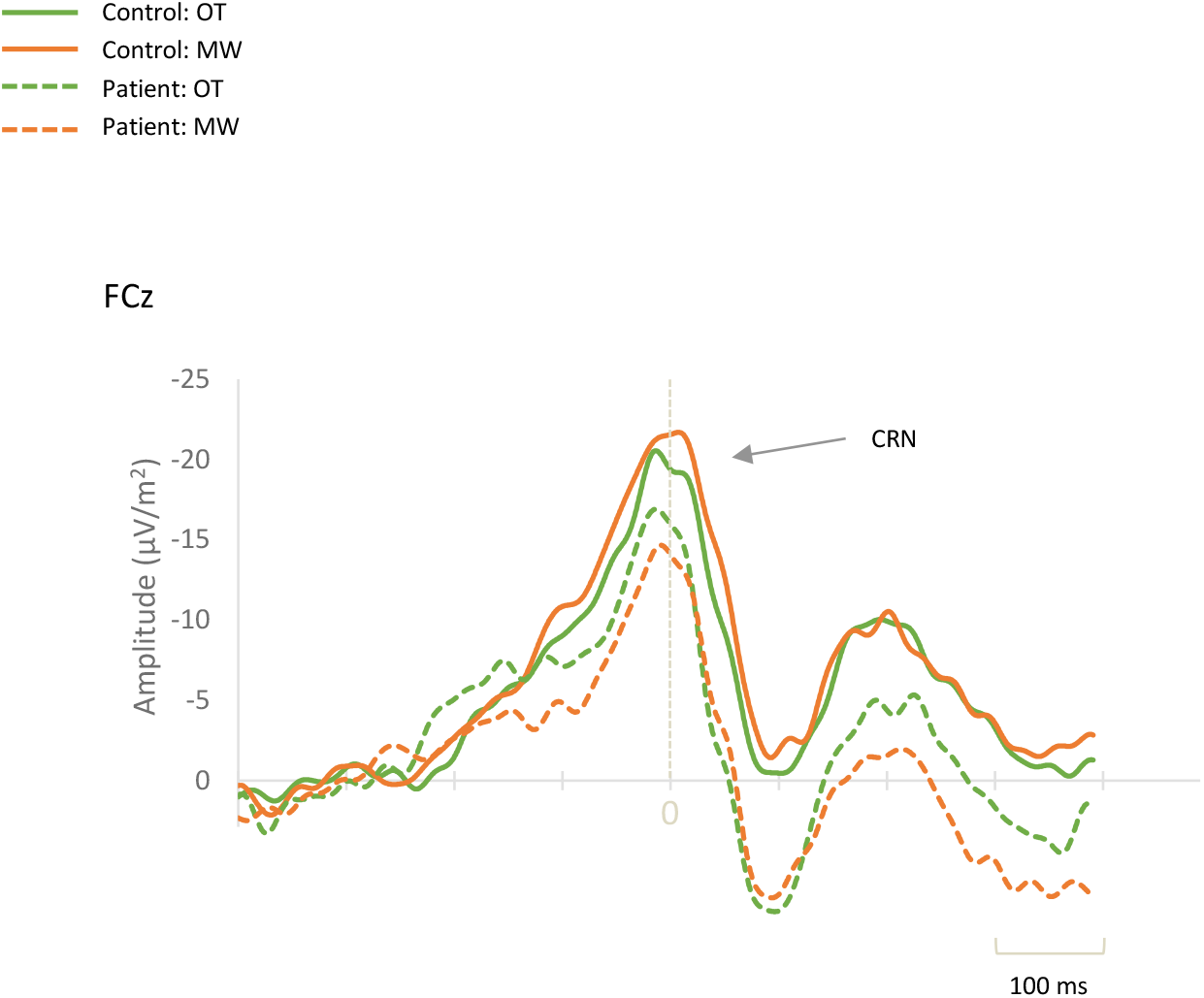
Grand average of the CRN response-related event-related potential (ERP) at the FCz electrode according to state (on-task [OT] or mind-wandering [MW]) and group (control or patient). CRN amplitude is significantly lower during MW than OT episodes in patients and significantly lower in patients than in controls specifically during MW episodes (*p* < 0.01).

## 4. Discussion

To specifically clarify the MW hypothesis (Bozhilova et al., 2018), our main goal in the present study is to confirm the negative impact of MW on perceptual and attentional processes while also exploring other cognitive processes, such as performance monitoring. The combination of thought-probe method and EEG recordings during a prolonged Go/No-Go SART confirmed that excessive MW-S in patients with ADHD was almost twice as frequent as in healthy controls. A differentiated negative impact of MW on performance was highlighted in both groups: MW affects RT speed and variability in patients and accuracy in controls. Finally, while perceptual and cognitive decoupling, which is commonly noted during MW, were observed in both groups, MW was associated with attenuated performance monitoring processes in ADHD patients exclusively, which provides valuable insight into cognitive mechanisms that contribute to MW-related impairments in ADHD.

First, regarding the phenomenological characteristics of MW, our results reveal that patients reported MW nearly twice as frequently (62%) as controls (32%), particularly in the case of MW-S, with a similar tendency observed for MW-D. To the best of our knowledge, the few studies that have used the thought-probe method to assess MW in patients with ADHD have exhibited a significantly higher MW frequency in patients, almost twice as much in some studies (Alperin et al., 2021; Bozhilova et al., 2021b; Gau et al., 2022; Madiouni et al., 2020). Moreover, distinguishing between the two types of MW advances our results a step further. Our findings indicate that MW-S is more strongly associated with the ADHD diagnostic itself than with a specific symptom, which is consistent with a clinical study that reports strong positive correlations between all ADHD symptoms and MW (described as less intentional) (Madiouni et al., 2020) as well as with a nonclinical study that indicates that all ADHD symptoms predicted MW-S (Arabacı and Parris, 2018). These results are also in accordance with an MRI study that was conducted with healthy controls in which researchers indicate that the MW-S tendency is negatively correlated with cortical thickness in the parietal and temporal regions and key regions of the DMN (Golchert et al., 2017). These same regions are often reported to have reduced cortical thickness in patients with ADHD (Liu et al., 2017; Makris et al., 2007; Narr et al., 2009). Second, our findings demonstrate that the frequency of MW-D is linked to the severity of impulsivity/hyperactivity symptoms. We must remain cautious on this point, as only 30% of the total MW frequency was deliberate, although this result is consistent with an MRI study that reveals, in healthy subjects, that cortical thickness in the frontal regions is specifically negatively correlated with MW-D (Golchert et al., 2017). Furthermore, altered activity as well as low cortical thickness in these regions is associated with higher impulsivity traits (Asahi et al., 2004; Lim et al., 2021; Russo et al., 2008; Shen et al., 2014). Our results, together with previous findings, suggest a complex relationship between ADHD symptoms and distinct neural correlates of MW-S and MW-D; consequently, it is crucial to examine the impact of both types of MW on performance and electrophysiological measures in ADHD, as they may be differentially affected. Finally, specifically regarding the subjective states assessed through the task, boredom, rather than motivation or fatigue, best predicts the MW state, whether spontaneous or deliberate. While a propensity for increased boredom in ADHD is well-documented in the literature (Golubchik et al., 2020; Malkovsky et al., 2012), our findings suggest, for the first time, that boredom is linked more with the MW frequency than the ADHD symptomatology or diagnosis. Notably, the elevated boredom levels observed in the ADHD group are significant only due to the increased frequency of MW in these patients because after adjusting for inter-individual differences of MW frequency, boredom levels were similar in both groups. Findings from recent fMRI studies have indicated that boredom and MW share overlapping neural correlates within the DMN (Danckert and Merrifield, 2018; Deng et al., 2022; Raffaelli et al., 2018); together with our results, these findings suggest that boredom and MW could be interrelated cognitive states that could reinforce each other.

In terms of performance, our main finding is that MW has a different negative impact for patients and controls. In comparison to OT episodes, accuracy was impaired during MW in controls (as revealed by a higher number of CE), while RT speed and variability were affected in patients with ADHD (as revealed by higher RT and *SD*-RT). These results, which are interesting for several reasons, offer insights into the sustained attention performance deficits seen in ADHD. First, the increase in CE during MW episodes has already been highlighted in controls (Chidharom and Bonnefond, 2023; Groot et al., 2021; Jin et al., 2019; Stawarczyk et al., 2014) and related to interference between the DMN and executive control networks (Christoff et al., 2009). In patients, however, our results suggest that CE are more closely associated with ADHD symptoms—specifically, impulsivity/hyperactivity—rather than with MW itself. Indeed, in patients, unlike in controls, MW had no impact on accuracy (they had similar numbers of CE during OT and MW episodes). This result may initially appear contradictory to that of Bozhilova et al., (2021), who report that MW negatively impacts accuracy only in ADHD patients; however, this discrepancy is likely due to differences in the nature of the tasks used. Bozhilova et al., (2021) used tasks that did not require participants to withhold a dominant motor response, which is unlike our task. Additionally, in our study, the high number of CE in patients, which significantly exceeded that of controls even during OT episodes, suggests a possible ceiling effect that may mask further performance declines during MW episodes. Patients’ overall higher number of CE during the entire task also supports this interpretation. Instead of the MW frequency, the impulsivity/hyperactivity score is the best CE predictor; these results align with the existing inhibitory deficits seen in ADHD patients (Fuermaier et al., 2015) and the overall altered FPN activity that is highlighted in ADHD (Gao et al., 2019). Second, and perhaps more interestingly, MW in patients with ADHD was characterized by slower and more variable RTs in comparison to OT episodes, and these two measures are known to be interrelated (Schmüser et al., 2016). Conversely, RT variability was not affected by MW in controls. The finding of higher attentional instability during MW episodes in patients suggests that this is the result of a combined effect of a trait (i.e., ADHD) and a state (i.e., MW) rather than a simple marker of the disorder. Accordingly, during OT episodes, as evidenced by comparable RTs and *SD*-RT in both groups, patients did not present greater attentional instability—in other words, a greater occurrence of moments of attentional disengagement—than controls, which aligns with study findings in which MW-S is a good predictor of speed impairments in ADHD patients (Bozhilova et al., 2020). In our study, identifying MW frequency as the best predictor of *SD*-RT, and determining a comparable overall RT between both groups throughout the task also supports our interpretation and MW hypothesis of ADHD proposed by Bozhilova et al., (2018), excessive MW in ADHD may contribute to performance impairments that classically describe this disorder.

Regarding our electrophysiological findings, perceptual and cognitive decoupling were observed during MW episodes in both groups. First, and in contrast to Bozhilova et al.’s (2021) findings, which suggest that poor adaptation of sensory processing is a primary deficit in ADHD, we find evidence of no such deficit in our study, which aligns with findings from other studies that exhibit intact P100 amplitudes in ADHD patients across various tasks (Kaiser et al., 2020; Peisch et al., 2021). However, our results are consistent with an overall attentional allocation deficit in ADHD (Peisch et al., 2021; Szuromi et al., 2011), as revealed by a lower P3b mean activity in patients. Then, we confirmed in both groups MW’s negative effect on sensory and attentional processing, which is widely described in the literature (Kam et al., 2022). This decoupling is typically attributed to the inverse relationship between the activation of the posterior cingulate cortex, which is a key DMN region, and the activity of primary sensory cortices and attentional networks (Bozhilova et al., 2018; Christoff, 2012; Christoff et al., 2016). In contradiction of certain assumptions of the MW hypothesis in ADHD (Bozhilova et al., 2018), however, our results do not reveal a more pronounced deleterious effect in patients during MW episodes. Rather, our results regarding performance monitoring processes allow us to progress in the initial hypothesis. To the best of our knowledge, and for the first time, we demonstrate a reduced performance monitoring ability specifically during MW episodes in ADHD patients, with no such impairment observed in controls. Supporting this finding, a recent study was conducted with healthy subjects and used the same task as our study, reveals reduced error monitoring, as measured by ERN amplitude, specifically during periods of suboptimal attention only in subjects with poor sustained attention abilities (Chidharom et al., 2021). In contrast, participants with good sustained attention abilities showed preserved performance monitoring — similar to the pattern observed in healthy controls of our study. Researchers who have, to date, reported impaired performance monitoring in ADHD patients, have not accounted for patients’ attentional state (Ehlis et al., 2018; Marquardt et al., 2018). Further studies are therefore needed to confirm the reduced efficiency of performance monitoring processes in ADHD patients during nonoptimal attentional states. Moreover (and compatibly with this first explanation), our results could suggest a particularly impaired level of arousal in patients during MW episodes. Indeed, a reduced performance monitoring ability, as revealed by reduced ERN amplitude and increased RT variability (as was the case in our study), have been evidenced in individuals with low brain arousal due to sleep deprivation (Hsieh et al., 2007; Renn and Cote, 2013; Tsai et al., 2005). This explanation would, in a way, reconcile some central postulates of the ADHD cognitive–energetic model, which emphasizes the role of arousal in ADHD performance deficits, with the MW hypothesis by suggesting a more direct interplay between hypo-arousal and the DMN interference (Hegerl and Hensch, 2014; Helfer et al., 2020; Isaac et al., 2024; Strauß et al., 2018).

We highlight MW experiences as excessively frequent in ADHD during an attentional task, which is associated with performance deficits that are typically described in this disorder; consequently, our results support the development of more targeted interventions to better regulate MW frequency in ADHD patients. From a mechanistic perspective, future studies should investigate the effects of methylphenidate treatment on performance monitoring processes to directly test its potential contribution to MW-related impairments in ADHD.

## Acknowledgments

This work was supported by the Jean-Marie Warter Prize 2017 and an internal grant funded by the University Hospitals of Strasbourg.

We thank the participants for their cooperation in this study.

