## Supplemental_information for "Effects of mind-wandering on cognitive and neural processes: Identifying specific impairments in adults with attention-deficit/hyperactivity disorder"

Supplementary methods

Statistical analysis

An ANOVA was performed on overall levels of different subjective states (motivation, boredom, and fatigue) between the two groups (control or patient). Then, an analysis of covariance (ANCOVA) was performed by adding mind-wandering (MW) frequency as a covariate. In order to verify that the frequency of MW was not driven by inter-individual differences in levels of motivation, boredom and fatigue, an ANCOVA on the frequency of MW was performed by adding each subjective state as a covariate. Sample *t*-tests were performed on overall behavioral measures (commission errors [CEs], reaction times [RTs], standard deviation of RTs [*SD*-RTs], efficiency score [ES]) to compare performance during the task between the two groups (control or patient). Additional analyses of correlation were performed between performance measures, the total MW frequency, and the three ASRS scores. Multiple regression analyses were performed to identify predictors of each performance measure (dependent variable) among the total MW frequency, the three ASRS scores, and group factor (independent variables). Means and *SD*s of overall performance and ERP measures during the whole task are presented in Supplementary table 1.

Supplementary results

The ANOVA performed on motivation, boredom, and fatigue revealed that each had a significant group effect: $F_{(1,54)} = 4.52$, $p < 0.05$; $F_{(1,54)} = 4.31$, $p < 0.05$; $F_{(1,54)} = 7.58$, $p < 0.05$, respectively. Motivation was significantly lower in patients (56.65 ± 24.18) than in controls (69.50 ± 20.95). Boredom and fatigue were significantly higher in patients (48.12 ± 24.98 and 54.76 ± 20.61, respectively) than in controls (34.25 ± 25.04 and 39.70 ± 20.30, respectively). After adding MW frequency as a covariate, the group effect was no longer significant for any of the subjective states ($p = 0.87$; $p = 0.69$; $p = 0.34$, respectively).

The ANCOVA performed on the frequency of MW revealed a significant group effect $F_{(1,51)} = 18.08$, $p < 0.001$. After controlling for inter-individual differences of levels of motivation, boredom and fatigue, the frequency of MW remained significantly higher in patients compared to controls.

The Mann–Whitney U-test performed on CEs revealed a trend effect of group (U = 279, $p = 0.06$). Patients tended to have a higher number of CEs than controls. The

Mann–Whitney U-tests performed on RTs and *SD*-RTs revealed no group effect (U = 362, $p = 0.63$ and U = 317, $p = 0.22$, respectively). The student's *t*-test performed on the ESs revealed a significantly lower ES in patients than in controls ($t_{(54)} = -2.74$, $p < 0.01$).

Correlation analysis revealed a significant positive link between the overall CEs and impulsivity/hyperactivity ($r_s = 0.35$, $p < 0.01$) and total ASRS scores ($r_s = 0.32$, $p < 0.05$) as well as a significant positive link between overall *SD*-RTs and MW episodes ($r_s = 0.31$, $p < 0.05$). A significant negative link was found between overall ESs and MW episodes ($r_s = -0.41$, $p < 0.01$) as well as between inattention and impulsivity/hyperactivity and total ASRS scores ($r_s = -0.26$, $p < 0.05$; $r_s = -0.32$, $p < 0.05$; $r_s = -0.31$, $p < 0.05$, respectively).

A multiple linear regression analysis revealed that the impulsivity/hyperactivity ASRS score ($\beta = 0.65$, $t = 3.25$, $p < 0.01$) is the predictor of CEs ($R^2 = 0.16$, $F_{(1,54)} = 10.6$, $p < 0.01$). The predictor of *SD*-RTs ($R^2 = 0.15$, $F_{(1,54)} = 9.57$, $p < 0.01$) and ES ($R^2 = 0.17$, $F_{(1,54)} = 11.6$, $p < 0.01$) is the MW frequency ($\beta = 6.85e-4$, $t = 3.09$, $p < 0.01$; $\beta = -0.55$, $t = -3.40$, $p < 0.001$, respectively) and the MW frequency ($\beta = 0.001$, $t = 2.86$, $p < 0.01$) and impulsivity/hyperactivity ASRS score ($\beta = -0.003$, $t = -2.80$, $p < 0.01$) are the predictors of RTs ($R^2 = 0.16$, $F_{(2,53)} = 5.26$, $p < 0.01$).

Supplementary table 1. Performance and event-related potential (ERP) measures during the whole task according to group (control or patient) (mean ± standard deviation).

| | Controls | | Patients | | | |
|---|---|---|---|---|---|---|
| | M | *SD* | M | *SD* | Test statistics | $p_{value}$ |
| CEs[a] | 18.85 | 10.65 | 27.85 | 16.81 | U=279 | 0.06 |
| RTs[a] | 0.40 | 0.05 | 0.40 | 0.10 | U=362 | 0.63 |
| *SD*-RTs[a] | 0.09 | 0.03 | 0.11 | 0.05 | U=317 | 0.22 |
| ES | 178.47 | 26.37 | 154.40 | 38.25 | $t_{(54)}$=-2.74 | < 0.01 |
| P100 (µV/m$^2$) | 34.32 | 17.45 | 37.54 | 20.53 | $t_{(49)}$=-0.60 | 0.55 |
| P3b (µV/m$^2$) | 18.72 | 9.49 | 12.46 | 7.73 | $t_{(49)}$=2.59 | < 0.05 |
| CRN (µV/m$^2$) | -21,53 | 10.88 | -17.24 | 9.06 | $t_{(49)}$=-1.66 | 0.10 |

Note. Student t-test; Mann-Whitney U-test; [a]Shapiro-Wilk is significant (p<0.05) indicating a deviation from normality

Abbreviations: CEs- commission errors, RTs- reaction times, *SD*-RTs- standard deviation of reaction times, ES- efficiency score
